# Quantitative CDK2 Dynamics Are Linked to Cell Fate Decisions in Differentiating Trophoblast Stem Cells

**DOI:** 10.64898/2026.05.17.725805

**Authors:** Sage Indigo G. Brill, Upasna Sharma, Estefania Sanchez-Vasquez, Ali Shariati

## Abstract

During early development of the placenta, a subset of murine trophectoderm stem cells (TSCs) undergo endoreplication, an unusual form of cell division cycle that decouples DNA synthesis from cytokinesis, resulting in physiological polyploidy. Oscillations in CDK2 activity are essential for the orderly progression of the cell cycle to ensure replicated DNA is accurately partitioned into two daughter cells. However, it remains underexplored how the dynamics of CDK2 activity regulate endoreplication in the context of TSCs differentiation. To address this question, we leveraged the variability in cell fate decisions in an established *in vitro* system of TSCs differentiation that relies on removal of a growth factor, FGF4, to induce endoreplication. Using quantitative single-cell live confocal microscopy of a precise CDK2 biosensor, DHB-Venus, we identified at least three different outcomes upon FG4 removal: self-renewal, endoreplication, and migration. Our quantitative analyses showed high levels of Cdk2 activity in self-renewing cells whereas intermediate DHB-Venus turnover is linked to increased nuclear and cell size, indicating a shift to endoreplication. Importantly, we also characterize a third class of differentiating TSCs with migratory characteristics that correlate with low levels Cdk2 activity without a change in nuclear size. In sum, our results demonstrated a correlation between different fate outcomes and specific thresholds of CDK2 activity. Our findings show that TSCs can distinguish between different outcomes through modulating the central kinase of the cell cycle, CDK2, positioning it as a key regulator of early trophoblast differentiation.

**Summary Statement:** This study investigates the oscillatory behavior of CDK2 activity during murine trophectoderm differentiation and its potential role in guiding cell fate decisions.

## Introduction

A typical mammalian cell division cycle allows cells to duplicate their DNA (S phase) and divide it between two daughter cells during the mitotic phase (M phase) (Mofatteh et al., 2021; Rubin et al., 2020). This canonical cell cycle is governed by oscillations in the activity of different cyclin-dependent kinases (CDKs) in complex with their specific activating cyclins (Herr et al., 2020; Liu et al., 2019; Sakaue-Sawano et al., 2013). The oscillatory activity of CDK2 in complex with Cyclin E (or Cyclin A) initiates the S phase, and the oscillatory activity of CDK1 in complex with Cyclin B drives the beginning of the M phase (Santamaría et al., 2007). However, not all cells in the body undergo mitosis after the S phase. A very distinct variant of the cell division cycle, termed endocycle or endoreplication, allows cells to skip the M phase to undergo multiple rounds of DNA synthesis, without cell division, to generate polyploid cells (Kołodziejczyk et al., 2023). Unlike polyploidy in cancer cells, developmental endoreplication is a programmed mode of cell division cycle that is linked to cell fate decisions through poorly defined mechanisms (Lee et al., 2009).

Mouse trophoblast stem cells (TSCs) are among canonical examples of cells that undergo developmentally controlled endoreplication to complete their differentiation program (Basak and Ain, 2022). During preimplantation embryonic development, diploid TSCs first emerge in the outer layer of a hollow spherical structure, known as the blastocyst, that also contains the inner cell mass, which gives rise to the embryo proper (Simmons et al., 2007). Mouse TSCs can undergo self-renewing division or differentiate into a multitude of cells, including polyploid trophoblast giant cells (TGCs) by undergoing endoreplication (El-Hashash et al., 2010; Hu and Cross, 2010). Successful endoreplication and differentiation to TGCs are essential for successful implantation in the endometrium (Woods et al., 2018). For instance, double deletion of E1 and E2 cyclins in mice results in defects in endoreplication of TSCs without affecting mitotic cells, leading to embryonic lethality due to placental defects (Parisi et al., 2003). Thus, it has been proposed that changes in the dynamics and threshold of CDK activity control endoreplication (Zielke et al., 2013). To test this hypothesis, we sought to test the relationship between quantitative variations in CDK2 activity and TSCs differentiation outcomes in a heterogeneous *in vitro* differentiation model.

Using a live fluorescent reporter, we monitored single-cell dynamics of CDK2 activity in TSCs as they underwent differentiation in response to FGF4 removal *in vitro* (Tanaka et al., 1998). Consistent with a recent report using a mouse model (Saykali et al., 2025), our findings show an overall decline in CDK2 activity during TSCs differentiation. Through long-term confocal imaging, we also uncovered how quantitative differences in the activity of the CDK2 are associated with self-renewal, endoreplication, and migratory behavior of TSCs. Our results further support the emerging evidence that links the dynamic behavior of CDKs activity with stem cell differentiation outcomes (Li and Kirschner, 2014; Ruijtenberg and van den Heuvel, 2016).

## Results

### Establishment of a dual biosensor system to monitor CDK2 activity in trophoblast stem cells

To investigate the cell cycle dynamics underlying early placental development, we first derived and established primary cultures of mouse TSCs from E3.5 blastocysts following previously published protocols (Seong and Rivron, 2023; Singh and Gerton, 2021; Tanaka et al., 1998).Immunofluorescence confirmed that the derived colonies robustly expressed the TSCs marker CDX2 (Niwa et al., 2005) (Supplementary Figure 1A). Upon induction of differentiation by FGF4 withdrawal for 48 hours, we observed undifferentiated TSCs that retained CDX2 expression, as well as the emergence of TGCs with no CDX2 expression (Quinn et al., 2006), suggesting a heterogeneous response to FGF4 removal (Supplementary Figure 1B).

To link CDK2 activity to TSCs differentiation outcomes, we engineered a dual fluorescent biosensor using a PiggyBac transposase-based vector to dynamically track the behavior of CDK2 during TSCs differentiation (Figure 1A). This construct expresses a previously validated CDK2 biosensor called DHB-Venus (Gu et al., 2004; Schwarz et al., 2018; Spencer et al., 2013) coupled with an mScarlet protein fused to a triple nuclear localization signal (3xNLS) for precise nuclear tracking (Figure 1A). This sensor design allows for the precise quantification of CDK2 activity based on cytoplasmic to nuclear localization ratio with higher ratio indicative of high Cdk2 activity. Upon establishment of the transgenic TSC line, we used image segmentation of the mScarlet signal to precisely quantify nuclear level of DHB-Venus, which were then dilated to measure the cytoplasmic level of DHB-Venus. Next, CDK2 activity was quantified as the ratio of the DHB-Venus mean intensity within this cytoplasmic shell to its intensity in the nucleus (Figure 1B). We observed distinct localization of the CDK2 biosensor in TSCs maintained in FGF4 compared to those undergoing FGF4-withdrawal differentiation, consistent with the hypothesis that quantitative changes in CDK2 activity are linked to TSCs differentiation (Figure 1C). Together, these results established a quantitative framework to monitor dynamics of CDK2 activity and directly link it to cell fate decisions during TSCs differentiation.

**Figure 1.**
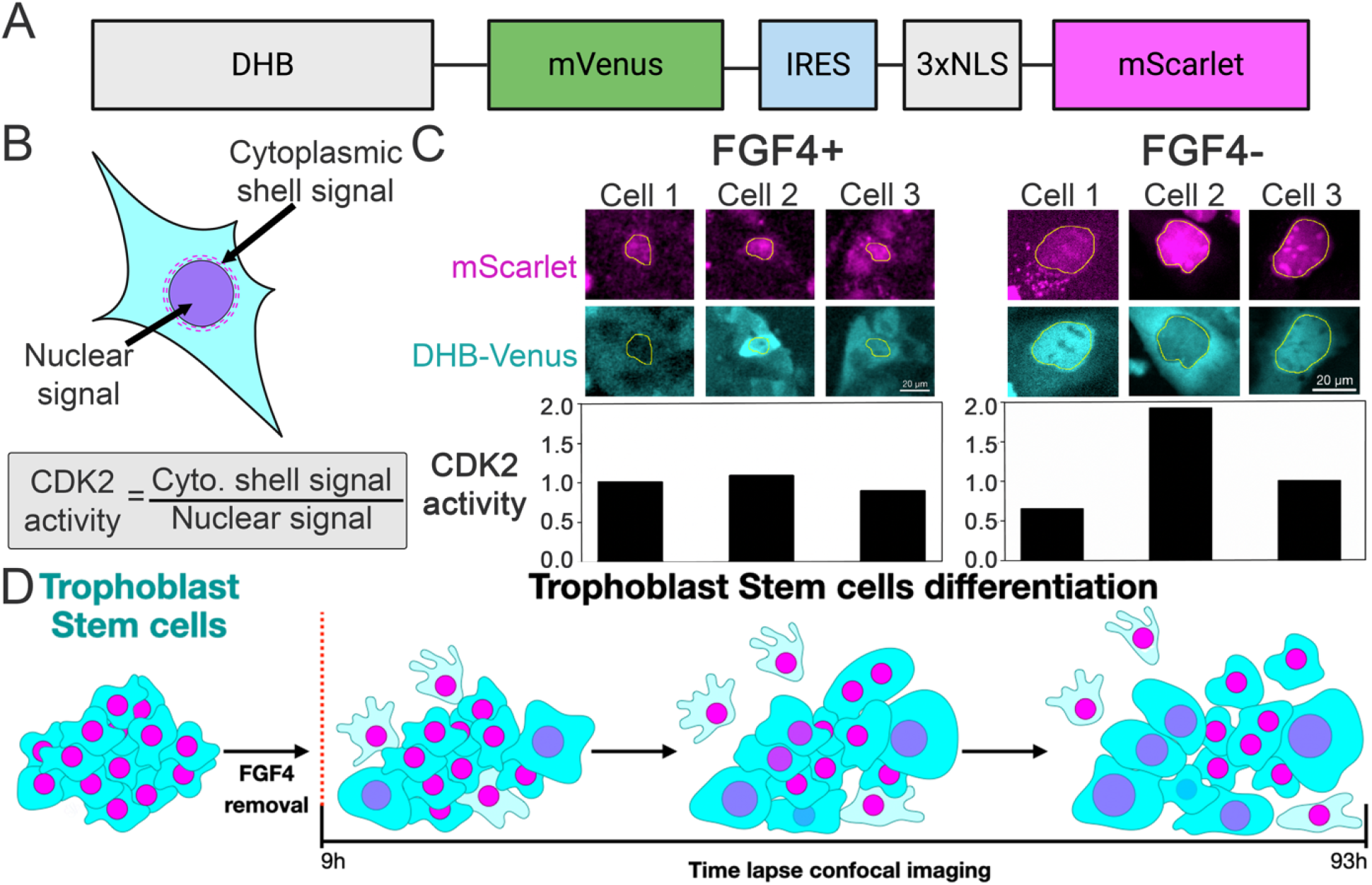
Optimization of a dual biosensor for CDK2 activity during *in vitro* differentiation of trophectoderm stem cells. Diagram of the dual biosensor utilized in this study for CDK2 activity quanhowification. The mRNA of the biosensor is formed by the amino acids 994–1087 of the human DNA helicase B (DHB) and the yellow fluorescent protein mVenus (DHB-Venus) linked via an IRES to a triple nuclear localization site (3xNLS) next to the mScarlet3 protein (mScarlet). This sensor enables nuclear segmentation and quantification of the cytoplasmic and nuclear DHB-Venus signal. (B) Advance segmentation was performed by using the mScarlet signal to create nuclear masks. Next, the cytoplasmic shell masks are a read between a 0.5µm and 2µm dilation of their specific nuclear masks to quantify the cytoplasmic shell signal. CDK2 activity is quantified as the ratio of the cytoplasmic shell mean intensity to the nuclear mean intensity of the DHB-Venus signal. (C) Representative images showing CDK2 activity in trophectoderm stem cells (TSCs) supplemented with FGF4 (FGF4+) and during TSCs differentiation, FGF4 withdrawal (FGF4-). (D) Schematic representation of the experimental design: confocal live imaging experiments on the differentiation of TSCs were performed for a period of seven days. Imaging started 9 hours after FGF4 removal.

### Live single-cell imaging reveals distinct cellular behaviors and CDK2 dynamics during TSCs differentiation

In order to monitor CDK2 dynamics during TSCs differentiation, we established a long-term confocal live-imaging pipeline using a spinning disk to reduce phototoxicity. We successfully extended our single-cell tracking window, capturing these dynamics for up to 93 hours, beginning 9 hours after FGF4 removal (Figure 1D), surpassing previous measurements (Saykali et al., 2025). By extending the duration of our live imaging experiments, we aimed to capture late-stage cellular behaviors during differentiation. Consistent with previous studies (Quinn et al., 2006; Saykali et al., 2025), we observed the emergence of cells undergoing self-renewing division and endoreplication with enlarged nuclei (Figure 2A). Furthermore, our analysis revealed a distinct subset of cells exhibiting migratory behavior, marked by active cellular displacement across the imaging field. Altogether, our qualitative analysis of the differentiating populations identified three distinct cellular trajectories: division, endoreplication, and migration (Supplementary Video 1 and 2).

**Figure 2.**
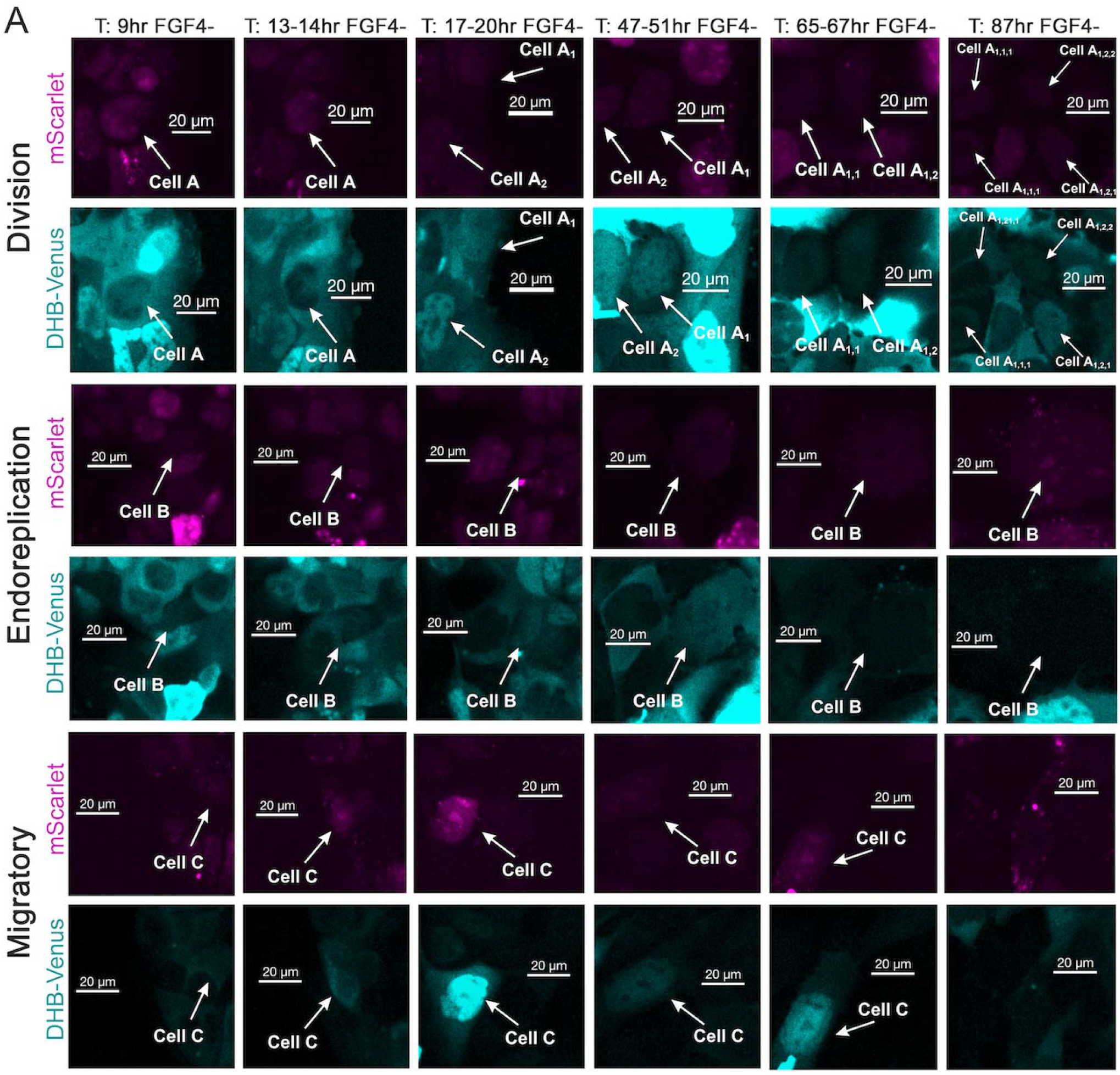
Confocal live imaging during the *in vitro* differentiation of trophectoderm stem cells identified several cellular behaviors. (A) Representative time-lapse images of differentiating trophectoderm stem cells expressing DHB-Venus and mScarlet. We classified the cellular behaviors of the differentiating trophectoderm stem cells following FGF4 removal as division, endoreplication, and migratory. Time points indicate hours post-FGF4 removal.

Next, we investigated whether the quantitative dynamics of CDK2 could distinguish between the different cellular outcomes during TSCs differentiation. From an initial dataset of approximately 300 imaged cells, we selected 60 individual cells for detailed downstream analysis based on strict data criteria: uninterrupted tracking fidelity over the seven-day period, a signal-to-noise ratio >5, and the absence of spatial overlap to ensure accurate segmentation. We classified these individual cells based on their CDK2 activity as measured by the cytoplasmic-to-nuclear ratio of the CDK2 reporter, with values above 1 indicating a predominantly cytoplasmic signal (high CDK2 activity), values around 1 indicating an even distribution between the cytoplasm and nucleus (intermediate CDK2 activity), and values below 1 indicating a primarily nuclear signal (low activity) (Supplementary Figure 2A). These data were plotted in a heatmap classifying differentiating cells into three major categories (Figure 3A): high CDK2 activity (ratio > 1.05), intermediate CDK2 activity (ratio 0.95–1.05) and low CDK2 activity (ratio < 0.95).

**Figure 3.**
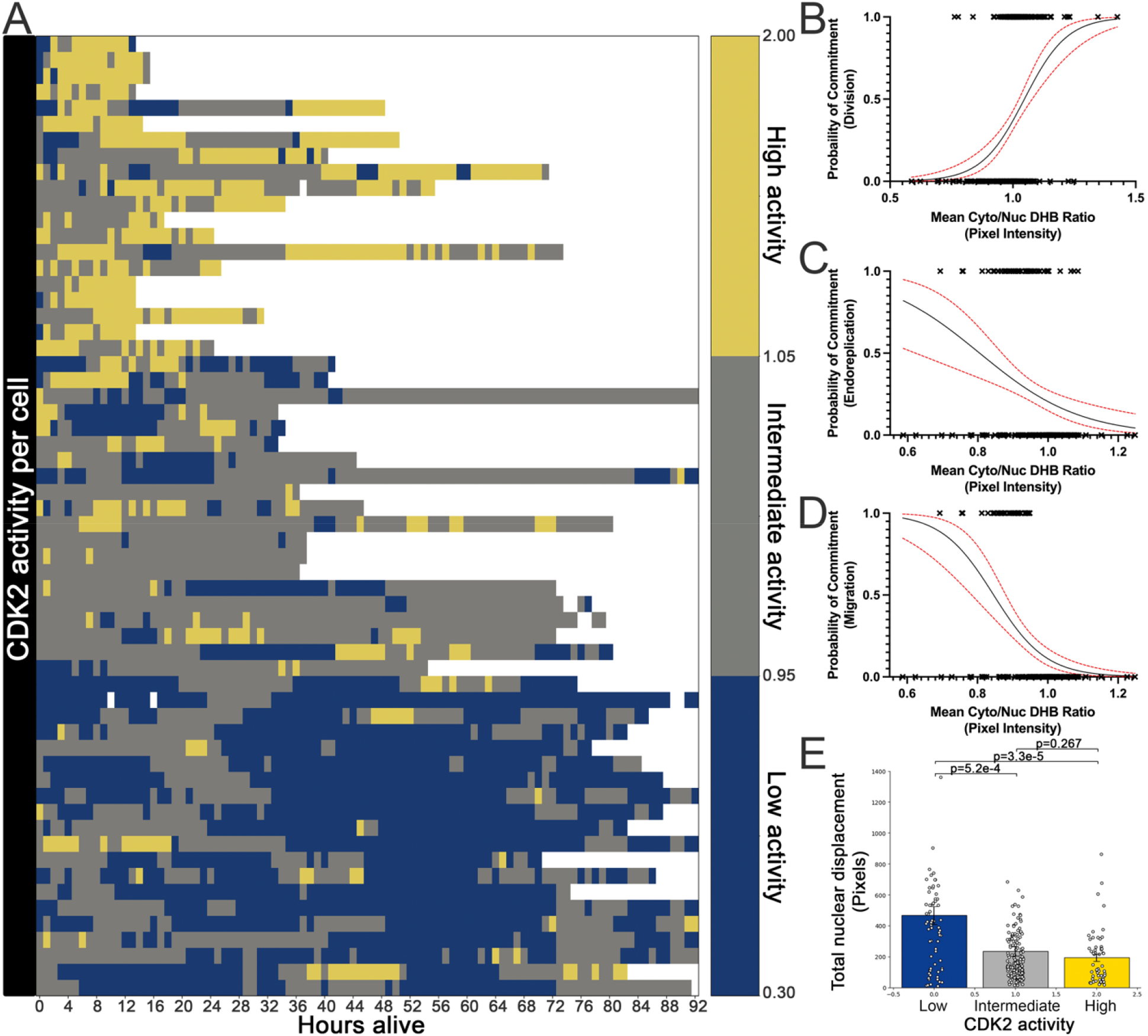
CDK2 activity is associated with cellular behavior during the *in vitro* differentiation of trophectoderm stem cells. (A) Heatmap of cytoplasmic-to-nuclear DHB-venus ratios over time for individual cells tracked following FGF4 removal. Cells are grouped by mean CDK2 activity: high (>1.05), intermediate (0.95–1.05), and low (<0.95). Time is aligned by hours alive per cell, with each daughter cell being treated as its own cell with time points. (B) Logistic regression predicts cell division commitment based on the mean cytoplasmic-to-nuclear DHB-Venus ratio. (C) Logistic regression prediction of migratory behavior based on the mean cytoplasmic-to-nuclear DHB-Venus ratio. (D) Logistic regression prediction of endoreplication commitment based on the mean cytoplasmic-to-nuclear DHB-Venus ratio. (E) Quantification of total nuclear displacement for non-dividing cells grouped by average CDK2 activity. Bars represent mean ± SEM. P values indicate Student’s t-test.

Logistic regression analysis showed that high CDK2 activity is predictive of TSCs undergoing cell division cycle, and to a lesser extent, low CDK2 activity was associated with migratory behaviors. However, intermediate CDK2 activity alone was not a strong predictor of endoreplication fate (Figure 3A-C). In addition, measuring the total nuclear displacement confirmed that cells with low CDK2 activity have significantly more cellular motility than those with intermediate or high CDK2 activity (Figure 3E). These analyses demonstrate that, during TSCs differentiation, quantitative changes in CDK2 ac ivity are correlated with distinct cellular fates.

### FGF4 withdrawal drives differentiation by modulating CDK2 activity

To measure the overall CDK2 activity during TSCs’ differentiation, we compared how the presence or absence of FGF4 impacts the activity of CDK2 over time. Our analysis demonstrated that, in the presence of FGF4, a significant proportion of cells (an average of 80% per frame) sustained high CDK2 activity, resulting in continuous self-renewal (Figure 4A). In agreement with previously reported data (Saykali et al., 2025), our analysis showed that upon induction of differentiation through FGF4 removal, the overall CDK2 activity progressively declined over time. This shift mostly gave rise to cells with an intermediate CDK2 activity that characterizes TGCs, averaging 38% per frame. Importantly, the proportion of cells with low CDK2 activity significantly increased from an average of 2.5% in the presence of FGF4 to an average of 25% per frame during differentiation. In addition, the removal of FGF4 was also associated with marked changes in nuclear area (Supplementary Figure 3A). Following FGF4 withdrawal, cells with intermediate CDK2 activity progressively increased their nuclear area without undergoing cell division (Supplementary Figure 3B). Meanwhile, cells with high CDK2 activity exhibited cyclic increases in the nuclear area over an average of 35 hours per cycle before undergoing division, while cells with low CDK2 activity associated with migratory behavior maintained a small nuclear area. This population-wide dampening of CDK2 activity and associated variations in nuclear size closely mirrored the phenotypic shift away from canonical cell division and toward increased rates of endoreplication and migration that characterized the first steps of murine placental differentiation (Jiang et al., 2023).

**Figure 4.**
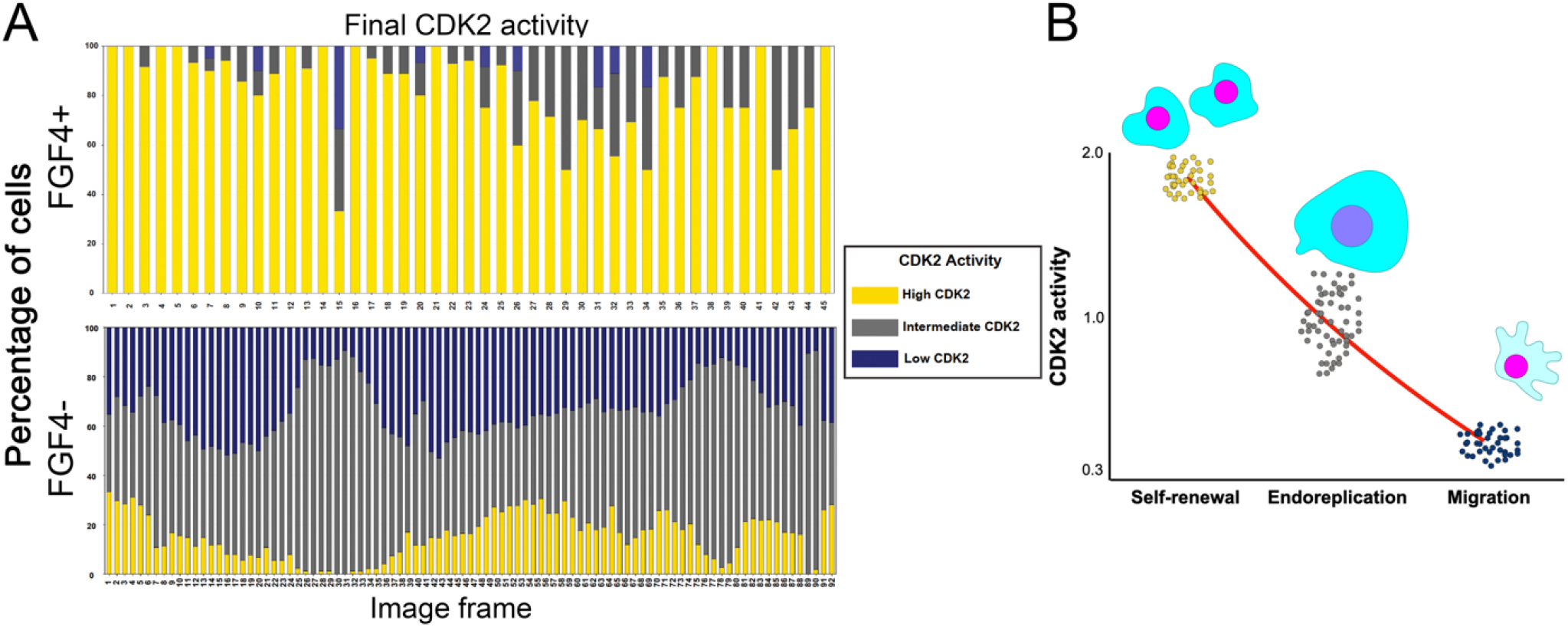
Removal of FGF4 during *in vitro* differentiation of trophectoderm stem cells changes the activity of CDK2 to allow migration and endoreplication. (A) Stacked bar plots representing the percentage composition of cells exhibiting high (yellow), intermediate (dark gray), or low (navy) CDK2 activity across time with and without FGF4. Upon removal of FGF4, CDK2 activity reduces, and as a consequence, the migratory and endoreplication behaviors increase. (B) Schematic illustration of our findings: CDK2 activity regulation is associated with different trophectoderm stem cell fate outcomes: division, endoreplication, or migration.

## Discussion

In this study, we establish that the quantitative dynamics of CDK2 activity are associated with cell fate outcomes in the murine trophectoderm. By deploying a live single-cell dual biosensor, we mapped the precise cellular trajectories of TSCs following the withdrawal of the stemness factor FGF4. As summarized in our working model (Figure 4B), we propose that different thresholds of CDK2 activity are associated with different TSCs differentiation outcomes. While sustained high CDK2 activity is essential for canonical cell division and self-renewal, the dampening of this kinase is a characteristic of differentiation. Specifically, an intermediate threshold of CDK2 activity is coupled with nuclear enlargement, steering cells toward endoreplication and the formation of polyploid trophoblast giant cells (TGCs). Conversely, cells with low CDK2 activity exhibit migratory behavior while maintaining a small nuclear size.

The placenta is an essential organ that supports nutrient exchange, hormone production, and immune regulation during embryonic development (Woods et al., 2018). The trophectoderm, which is the precursor to the placenta, is made up of trophectoderm stem cells (TSCs) that can be used to create *in vitro* models of differentiation to trophoblast giant cells (TGCs) (Quinn et al., 2006). Our data show that the proportion of endoreplication-like cells indeed increases upon FGF4 removal, which is consistent with previously described transitions along the TSC-to-TGC differentiation pathway (Tanaka et al., 1998). Nevertheless, during normal *in vivo* development, the establishment of a functional maternal-fetal interface relies heavily on the profound invasive and migratory capacities of specific trophoblast lineages (Aplin and Ruane, 2017). A recent high-resolution single-cell roadmap of murine placentation revealed that migratory trophoblast derivatives are associated with TGCs development (Jiang et al., 2023). Aligned with these findings, our data demonstrate that a distinct, non-dividing population of migratory cells also arises *in vitro* upon FGF4 removal. The identification of this highly motile, low-CDK2 population, has implications for the fidelity of placental differentiation models, as these cells likely represent the *in vitro* equivalent of migratory *in vivo* populations. Altogether our results suggest that the sharp down-regulation of CDK2 activity is linked to TSCs differentiation, likely through cell cycle and cytoskeletal remodeling required for successful trophoblast migration and endoreplication.

CDK2 is a known central regulator of the cell cycle, functioning primarily in complex with cyclins E and A to promote the transition to DNA synthesis in the canonical mitotic cell cycle. During the transition to endoreplication, mitotic entry is actively suppressed through the inhibition of CDK1, allowing cells to bypass mitosis and instead undergo repeated rounds of DNA synthesis (Fox and Duronio, 2013). Consequently, TSCs primarily rely on the regulated activity of CDK2 and its downstream E2F family transcription factors to undergo successive rounds of DNA replication without mitotic division (Chen et al., 2012). Our findings complement recent discoveries of an overall decline in CDK2 activity upon mouse embryo *in vitro* implantation (Saykali et al., 2025). However, our single-cell regression models uncover that the magnitude of this decline is what ultimately dictates specific lineage commitments: intermediate and low CDK2 states are uniquely associated with endoreplication and migration, respectively. Importantly, in normal placental development, CDK2 activity must be tightly regulated, as aberrant increases in its activity promote pathologic outcomes, such as trophectoderm-derived tumors (Olvera et al., 2001), likely due to the inappropriate maintenance of stemness in the placenta.

While we provide correlative evidence between CDK2 activity levels and cell fate outcomes during TSCs differentiations, further studies are needed to determine whether different CDK2 activities are causally linked to cell fate decisions through phosphorylation of distinct substrate groups. Considering that impaired trophoblast giant cell development and defective trophoblast invasion are hallmark features of various placental diseases (Cabunac et al., 2021; Knöfler et al., 2019; Ma et al., 2023; Pandey et al., 2023), deciphering the molecular drivers of these fate decisions is of clinical significance. Our findings determined how cell cycle dynamics intersect with lineage commitment in the placenta and position the regulation of CDK2 activity as a fundamental step for mammalian trophectoderm differentiation, offering new avenues for investigating the cellular origins of implantation dysfunctions.

## Materials and Methods

### Animals

Wild-type B6SJLF1/J mice were obtained from JAX Laboratories (Strain #: 100012). All animal procedures were approved by the UCSC Institutional Animal Care and Use Committee (IACUC) under protocol Sharu2410-01 (PI: Upasna Sharma). Mice were group-housed (maximum of 5 per cage) with a 12-h light-dark cycle (lights off at 6 p.m.) and free access to food and water *ad libitum*. Natural mating was performed to obtain E3.5 embryos for subsequent experiments.

### Trophectoderm stem cells derivation

We derived trophectoderm stem cells (TSCs) from E3.5 B6SJLF1/J blastocysts following previously published protocols (Seong and Rivron, 2023; Singh and Gerton, 2021; Tanaka et al., 1998). We used mitomycin-C-treated B6SJLF1/J mouse embryonic fibroblasts (MEFs), derived in the lab, for blastocyst outgrowth. Blastocysts were cultured on MEFs in TSCs derivation medium (RPMI 1640, 20% FBS, FGF4, heparin, sodium pyruvate, Glutamax, 2-mercaptoethanol, pen/strep). After outgrowth formation (∼3–5 days), colonies were disaggregated using TrypLE and passaged onto fresh MEFs. Following culture with media containing heparin (Sigma Aldrich # 501657295, 1mg/mL - 1000x) and FGF4 (R&D # 7460F4025, 25µg/mL-1000x), we picked TSCs colonies for further MEF-free culture. Early passage TSCs (< passage 20) were used for all experiments to preserve self-renewal potential.

### Trophectoderm stem cells maintenance

Established B6SJLF1/J TSCs were cultured in MEF-free conditions by maintaining them in TSC medium (RPMI 1640, 20% heat-inactivated FBS, FGF4 (25 µg/mL stock), heparin (1mg/mL stock), 2-mercaptoethanol, sodium pyruvate, Glutamax, penicillin/streptomycin, and 70% of MEF-conditioned medium), with the media being refreshed every 48 hours (Singh et al., 2023). TSCs were frozen in TSC freezing medium (TSC medium, 50% FBS, 20% DMSO) and stored in liquid nitrogen.

### Trophectoderm stem cells differentiation

To induce differentiation, TSCs were cultured in TSC medium lacking FGF4 and heparin for up to 8 days (Basak and Ain, 2022; Singh et al., 2023). Media refresh happened every 48 hours. Differentiation was monitored by cell morphology and nuclear size. TSCs seeding depended on vessel size: 8-well ibidi chamber -> 5 × 10^3^ cells/well, 12-well plate -> 1 × 10□ cells/well, 35 mm dish -> 2.5 × 10□ cells/dish. Cells were grown until ∼50% confluency before differentiation.

### Immunofluorescence and confocal microscopy

TSCs were cultured in µ-Slide 8 Well ibiTreat chambers (Ibidi USA, 80826) until colony formation. Next, we fixed the samples in 3.6% PFA for 10 minutes at room temperature. Cells were washed three times with TBS with 0.1% Tween-20 (TBSt). Blocking was performed with PBS containing 3% BSA (Thermo Fisher Scientific, 15260037) for 2–3 hours. The primary antibody rabbit monoclonal anti-CDX2 (D11D10) (Cell Signalling Technology, 12306) was diluted at 1:500 in blocking buffer and incubated overnight at 4°C. The following day, TSCs were washed with TBSt three times and incubated with the following secondary antibodies: Goat anti-Rabbit Alexa Fluor-647 (Thermo Fisher Scientific, A31556) and Alexa Fluor™ Plus 405 Phalloidin (Thermo Fisher Scientific, A30104) at 1:1000 dilution in blocking buffer for 1 hour. Following the secondary antibodies, TSCs were washed three times with TBSt before incubating for 30 min with 1:500 DAPI. Final washes were performed three times with PBS before imaging. Confocal microscopy was performed with an Oxford Instruments Andor BC-43, 20x dry objective NA 0.8.

### Construct design

To monitor CDK2 activity during TSC differentiation, we used fusion PCR to construct a dual fluorescent biosensor based on the DHB domain of the human DNA helicase B gene (https://www.addgene.org/136461/) coupled by an IRES to 3xNLS (nuclear localization sites)-mScarlet, a bright monomeric fluorescent protein that localizes to the nucleus (https://www.addgene.org/189775/). This dual reporter was cloned into a PiggyBac (PB) transposon vector for transfection (https://www.addgene.org/219794/). This design allowed the expression of DHB-venus and 3xNLS-mScarlet under the cytomegalovirus (CMV) promoter. In parallel, the backbone includes a puromycin resistance gene under the hPGK promoter for antibiotic selection. The final plasmid: PB-DHB-Venus-3xNLS-mScarlet (PB:DHB-Venus/Scarlet) was verified by Sanger sequencing to confirm sequence fidelity.

### Transfection and selection

For integrating PB:DHB-Venus/Scarlet into TSCs, we used Lipofectamine 3000 (Thermo Fisher, L3000001), diluted in OptiMEM (Thermo Fisher, 31985062) according to the manufacturer’s instructions. Briefly, TSCs were seeded in 0.1% gelatin-coated 6-well plates as described previously. The next day, cells were transfected in OptiMEM as followed: 200ng PiggyBac Transposase (SBI, PB210PA-1), 600ng PB:DHB-Venus/Scarlet and 4uL of Lipofectamine. Media was replaced 24 hours after transfection, and fluorescence was checked for transfection efficiency. Selection with 0.5ug/ml Puromycin started 48 hours after transfection and continued until nontransfected control cells died. Furthermore, Venus and mScarlet fluorescence were used to confirm stable integration.

### Live imaging experiments

For live imaging experiments, TSCs were cultured in µ-plate 24 Well ibiTreat (Ibidi USA, 82406) until colony formation, ∼3-4 days. We used a spinning disk confocal microscope, the Andor BC-43, equipped with a built-in incubator maintained at 37°C, 5% CO_2_, 20% O_2_, and 80% relative humidity. Imaging was performed with a 20x dry lens and 488 and 561 lasers to capture images of Venus and mScarlet, respectively. <u>Single-cell live imaging of the differentiation process</u> was initiated 9 hours after the removal of FG4 and heparin. Thirty TSCs colonies were selected based on the following criteria: absence of endoreplicating cells in the surrounding area, colony size ≤ 40 cells, strong DHB reporter signal in confocal channels, and central positioning within the well (avoiding edge aggregation). Automated time-lapse imaging was performed at 1-hour intervals over 93 hours. <u>Single-cell live imaging of undifferentiated TSCs</u> was initiated 48 hours after seeding in media with FGF4 and heparin. Fourteen TSCs colonies were selected based on the following criteria: absence of endoreplicating cells in the surrounding area, colony size ≤ 20 cells, strong DHB reporter signal in confocal channels, and central positioning within the well (avoiding edge aggregation). Automated time-lapse imaging was performed at 1-hour intervals over 45 hours.

### Image preprocessing and quality control

Preprocessing was performed on each subsection of cells, consisting of stitching images together to create a continuous string of photos, then applying a Gaussian filter to each image. DHB-Venus channel filtering was set to 0.312 µm, and mScarlet channel filtering to 0.250 µm. Each subpopulation was then analyzed across the full set of images, with analysis viability determined by evaluating for instances of lost signal and overall intensity of both nuclear mScarlet and DHB-Venus signals. For quality control, a qualitative review was performed to select sections of data with the fewest visual dark spots, ensuring smooth data analysis, and to confirm the presence of a consistent nuclear signal and measurable DHB-Venus expression.

### Image analysis

Time-lapse data was loaded into Fiji (Schindelin et al., 2012) for advanced segmentation and fluorescent quantifications. The mScarlet signal was used for advanced nuclear segmentation, which created multiple regions of interest (ROIs) via the freehand loop tool. The accuracy of the created ROIs was checked by overlaying them on the DHB-Venus signal and comparing them to the original full image. To study CDK2 activity, we quantified the ratio of DHB-Venus signal in the cytoplasm vs. the nucleus, following published protocols (Spencer et al., 2013). First, we made masks for the cytoplasm by expanding the initial outlines of the nuclei by a small amount measured in micrometers: each nuclear ROI was dilated by both 0.5 µm and 2 µm. Consequently, we applied an XOR function between the original and dilated ROIs, generating cytoplasmic shells that excluded nuclear regions. Nuclear ROIs and cytoplasmic shells were then separately masked over the isolated DHB-Venus channel, and mean intensity measurements were exported as .csv files for each cell. In sum, CDK2 activity = Cytoplasmic Mean intensity value / Nuclear Mean intensity value. The values of the above equation were used to classify cells based on DHB-Venus intensity, with values above 1 indicating predominantly cytoplasmic, values around 1 indicating an even distribution between cytoplasm and nucleus, and values below 1 indicating primarily nuclear intensity. The resulting ratios were analyzed to qualitatively group cells by high, intermediate, and low CDK2 activity and associate them with three common cell fate behaviors: division, endoreplication, and migration. In the case of cell division, each daughter cell was treated and normalized as its own individual cell lineage. Mean intensity ratios were plotted over time using the matplotlib library in Python.

## Acknowledgements

We gratefully acknowledge the support of the Department of Biomolecular Engineering (BME) at the University of California, Santa Cruz (UCSC). We extend our sincere thanks to the directors of the Institute for the Biology of Stem Cells (IBSC), Dr. Lindsay Hinck and Dr. Camilla Forsberg, for their continued leadership and support of this research community.

## Author Contribution

ASH and ESV conceptualized the project. ESV and ASH wrote the manuscript. SB and ESV performed all experiments, analyzed the results, and visualized the data. US contributed to the establishment of TSCs culture.

## Supplementary Figures

**Supplementary Figure 1.**
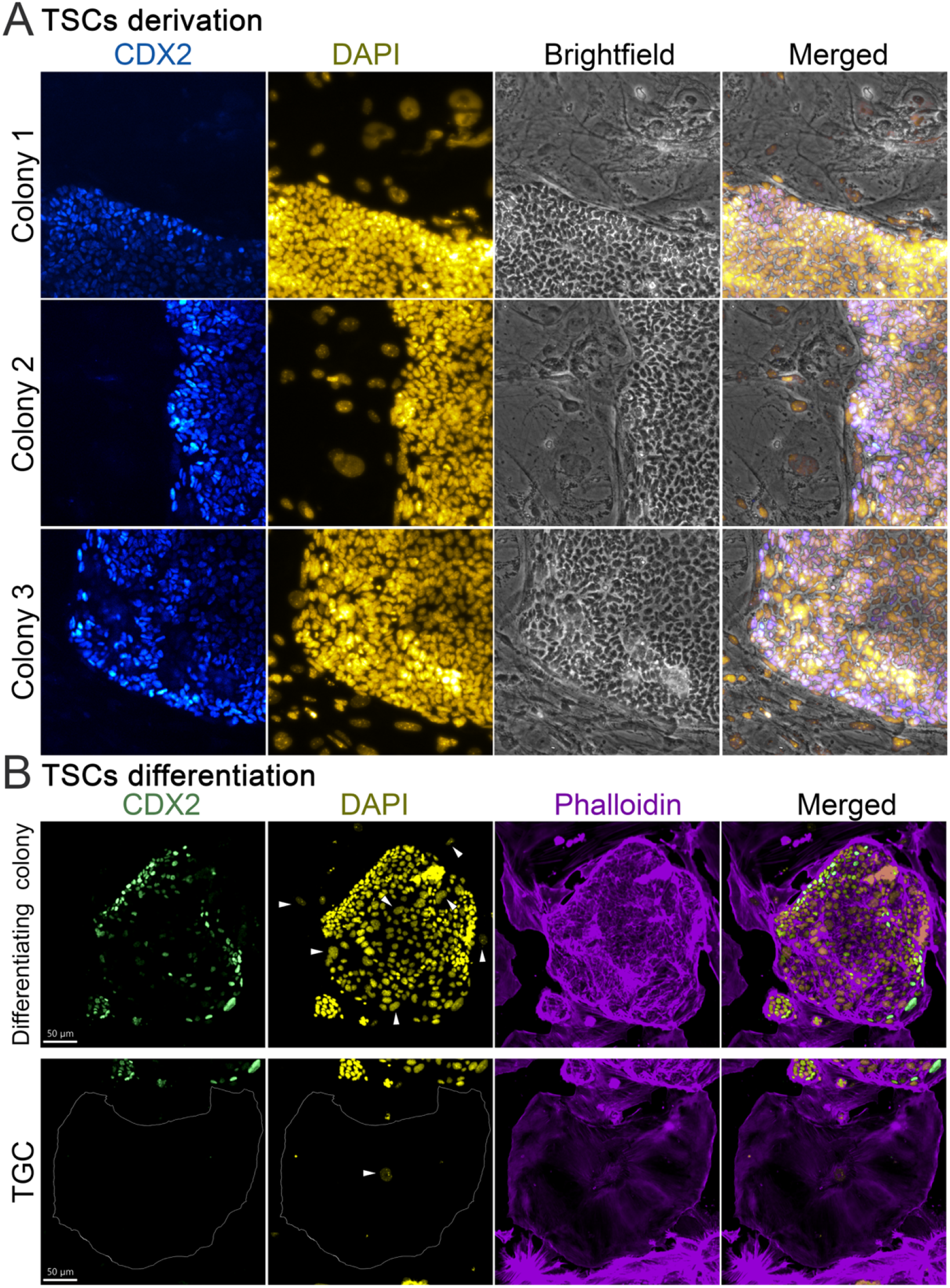
Trophectoderm stem cells derivation and differentiation. (A) Representative trophectoderm stem cells (TSCs) colonies immunostained with the specific marker CDX2 (blue). Nuclei were stained with DAPI (yellow). (B) After 48 hours of FGF4 removal, some differentiating TSCs maintain their colony morphology and have some CDX2 (green) positive cells. In parallel, some TSCs differentiate into trophoblast giant cells (TGCs), characterized by being CDX2 negative. Representative TGCs nuclei are marked by arrowheads. Nuclei were stained with DAPI (yellow) and membranes with phalloidin (violet).

**Supplementary Figure 2.**
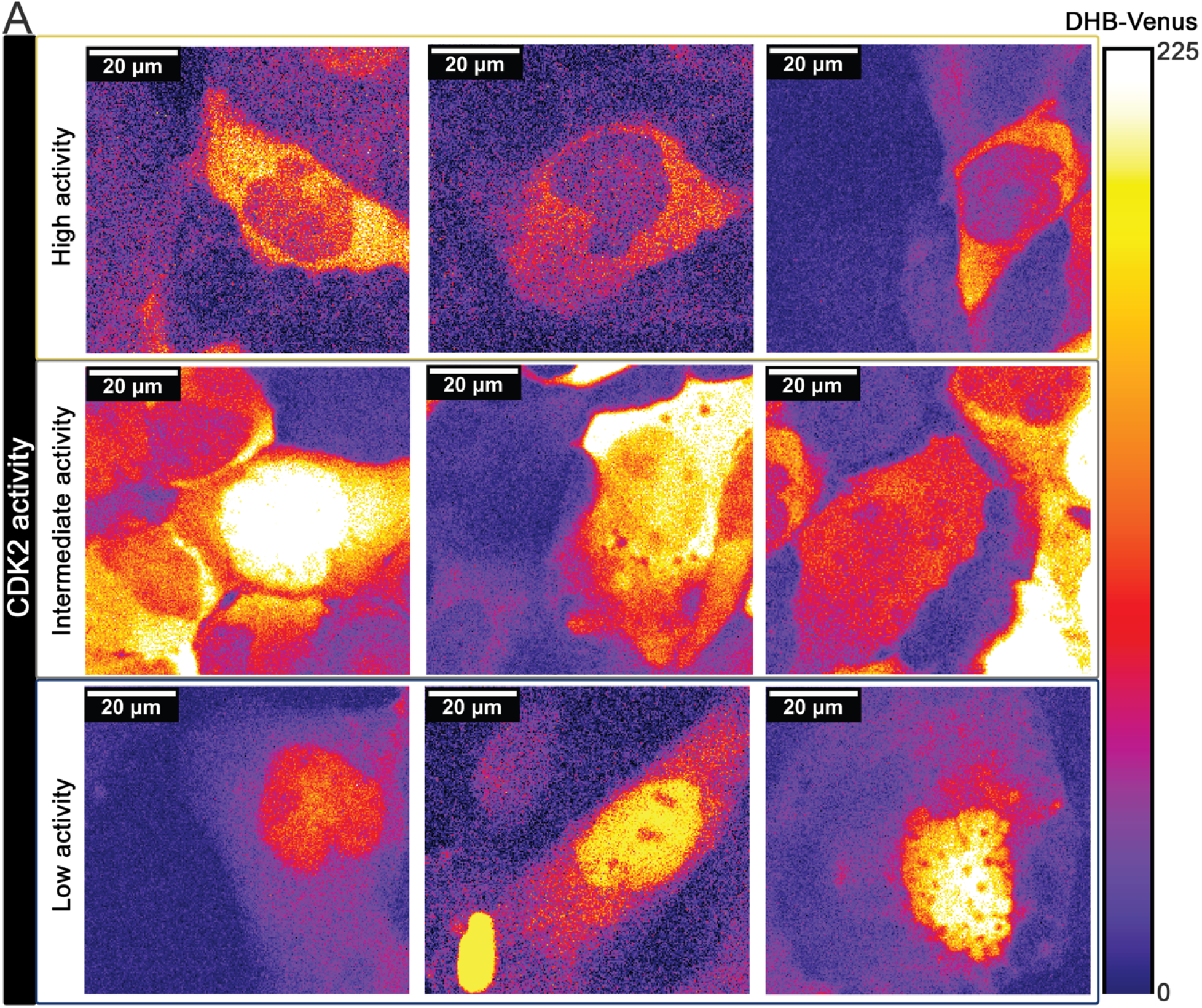
Classification of differentiating trophectoderm stem cells based on their CDK2 activity. (A) Representative images of differentiating trophectoderm stem cells classified according to their CDK2 activity: cytoplasmic-to-nuclear DHB-Venus ratio. Images are false-colored to visualize DHB-Venus intensity’s distribution. The color bar to the right reflects the pixel intensity range from 0 to 225 arbitrary units (A.U.)

**Supplementary Figure 3.**
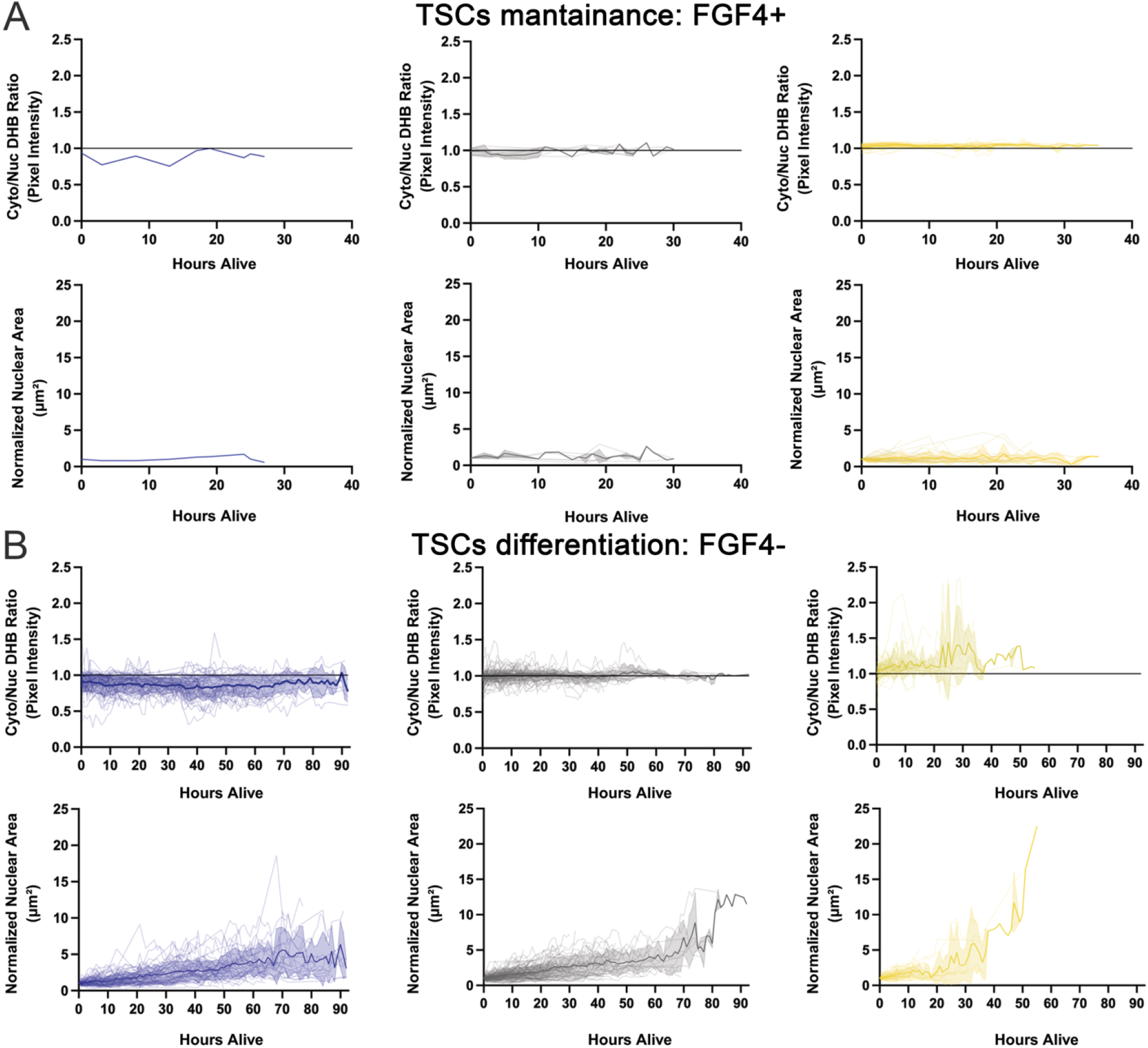
Arrays of behaviors during *in vitro* differentiation of trophectoderm stem cells are associated with diverse CDK2 activity. The changes in the ratio of cytoplasmic to nuclear DHB-Venus over time are shown for each group: division, endoreplication, and migration, with FGF4 (FGF4+, A) or without (FGF-, B). Smoothed mean trajectories with SEM shading reflect typical CDK2 activity dynamics.

**Video 1. Representative example of a differentiating trophectoderm stem cells (TSCs) colony selected for further analysis**. The video depicts a target TSCs colony meeting the strict morphological and spatial inclusion criteria established for downstream single-cell tracking. To ensure accurate quantification and tracking fidelity, selected colonies were required to demonstrate a size of ≤ 40 cells, an absence of multiple giant cells in the immediate surrounding area, central positioning within the well to avoid edge-aggregation artifacts, and a robust DHB-Venus reporter signal across the appropriate confocal channels.

**Video 2. Single-cell tracking of distinct cellular behaviors during trophectoderm differentiation**. Time-lapse spinning-disk confocal imaging of cells expressing the CDK2 biosensor (DHB-Venus) and nuclear marker (mScarlet) over a seven-day period. The video highlights representative examples of the three cellular behaviors quantified in the study. A blue tracked dot with a continuous line illustrates a single cell exhibiting migratory behavior. A solitary blue dot denotes a cell undergoing endoreplication. White arrowheads follow cells undergoing canonical cell division.

